# Neural and behavioral dynamics of encoding, production and synchronization with external rhythms in subcortical lesion patients

**DOI:** 10.1101/2024.01.06.574472

**Authors:** Antonio Criscuolo, Michael Schwartze, Sylvie Nozaradan, Sonja A. Kotz

## Abstract

Acting in and adapting to a dynamically changing environment necessitates precise encoding of the *timing* of unfolding sensory events in our environment, and to *time* of our own (re-)actions to them. Cerebellar (CE) and basal ganglia (BG) circuitries play fundamental and complementary roles in timing processes. While the CE seems to encode the precise timing of sensory events (*when* an event occurs), the BG engage in generating temporal predictions (*when* a next event occurs). However, their contributions are rarely investigated in combination, as it is generally difficult to record data from respective patient groups in parallel.

Here we investigated the causal roles of CE and BG in sensory and sensorimotor timing processes. Healthy controls and patients with CE or BG lesions listened to isochronous auditory sequences while their EEG was recorded and later performed a tapping synchronization task. We assessed intra- and inter-individual variabilities, as well as group differences, using event-related responses, delta-band inter-trial phase-coherence and acceleration dynamics while tuning to the stimulation frequency (*Sf*). CE and BG lesions increased variability in ERP latency and reduced the coherence of delta-band activity. CE but not BG lesions further impacted the stability of delta-band oscillations while tuning to the *Sf*. These findings show a causal link between subcortical lesions and the capacity to encode and synchronize ongoing neural activity with temporal regularities in the acoustic environment, but do not fully dissociate the specific contributions of the BG and the CE to processing sound in isochronous contexts.

## 1. Introduction

Time is a fundamental dimension of human cognition. Every decision, every action, every sensory stimulus around us happens in time and at multiple timescales. Our capacities to encode the precise *timing* of events and to *time* our (re-)actions to them are pivotal for acting in and adapting to a continuously changing dynamic environment. However, the environment displays some gradient of *temporal regularity* (Greenfield, Honing, et al., 2021): there are various periodicities within the body (e.g., the heartbeat and respiration; Criscuolo et al., 2022) as well as in simple and complex sensory input (e.g., music, speech) and in behavior (e.g., interpersonal interactions in the animal and human world; Greenfield, Aihara, et al., 2021). These temporal regularities (or *rhythm*) can facilitate the processing of the *timing* of sensory events, and further enable individuals to anticipate the *when* of next sensory events. In turn, temporal anticipation can benefit *adaptive* behavior: we can dance to music because we know *when* the next beat falls, and we can synchronize with others because their movement timing is predictable. Yet what determines our capacity to adapt behavior in time? Do we differ in the *if* and *how* we adapt and synchronize with auditory rhythms?

We have previously proposed (Criscuolo et al., 2023) that synchronization depends on two fundamental lower-level capacities that form a 3-node framework: detect, produce, and synchronize (‘DPS’) with rhythms (Criscuolo et al., 2023). Encoding is a prerequisite to detect rhythms. Human (e.g., Schroeder & Lakatos, 2009) as well as nonhuman animals (e.g., Lakatos et al., 2008) can do so by aligning endogenous neural oscillatory activity at multiple timescales (Criscuolo et al., 2023; Lakatos et al., 2005) to temporal regularities in the sensory environment. Detecting regularities allows generating predictions about *when* next events occur, thus enabling *dynamic attending* (Large & Jones, 1999) of event onsets. Temporal predictions materialize as anticipatory neural activity (Arnal, 2012; Breska & Deouell, 2017a; Fujioka et al., 2009, 2012, 2015; Ross et al., 2018; Snyder & Large, 2005), which supports the predictive alignment of neural oscillations to event onsets, fosters sensory processing (Lakatos et al., 2013) and further allows for production and synchronization of behavior in a predictive manner (Bartolo & Merchant, 2015; Gámez et al., 2018). Thus, “adaptation by anticipation” (Fraisse, 1963; p. 18) is a mechanism for optimizing behavior and operates across sensory modalities (Arnal & Giraud, 2012; Cravo et al., 2013; Friston, 2005; Morillon et al., 2016; Nobre et al., 2012).

Fundamental to predictive temporal processes is the engagement of an extended motor system (Arnal, 2012; Fujioka et al., 2012, 2015; Schwartze & Kotz, 2013), recruited to prepare and execute predictive and synchronized behavior. The cerebellum (CE) and the basal ganglia (BG) are structures in an extended subcortico-cortical brain network that plays a pivotal role in temporal processing (Merchant, Harrington, et al., 2013b; Schwartze & Kotz, 2013). The CE encodes the precise timing (*when* an event onsets) of sensory events in the subsecond range (Bareš et al., 2018; Ivry et al., 1988; Ivry & Keele, 1989; Ivry & Schlerf, 2008) and thus allows estimating the duration of temporal intervals (Breska & Ivry, 2020, 2021; Grube, Cooper, et al., 2010; Grube, Lee, et al., 2010; Teki, Grube, & Griffiths, 2011; Teki, Grube, Kumar, et al., 2011) by quantifying the time elapsed between event onsets. The BG support the generation of temporal predictions (*when* the *next* event occurs) and use relative timing to process the beat (Grahn, 2009; Grahn & Brett, 2009a; Schwartze et al., 2011a; Teki, Grube, Kumar, et al., 2011). Consequently, patients with BG lesions are less sensitive to temporal regularity in auditory sequences, potentially resulting in a less efficient prediction of incoming sensory information in basic (Schwartze et al., 2015) as well as in complex (syncopated rhythms; Nozaradan et al., 2017) sequences. In contrast, CE lesions tend to not impact the capacity to generate temporal predictions (Schwartze et al., 2016b) but alter the encoding of event onsets in basic and complex sound sequences (Nozaradan et al., 2017), ultimately resulting in delayed and variable early event-related responses in the EEG (Schwartze & Kotz, 2021). Altered encoding of event onsets affects the capacity to encode and predict temporal intervals (Breska & Ivry, 2018, 2020, 2021; Grube, Cooper, et al., 2010) and further impacts the ability to produce and synchronize with rhythms (Ivry & Keele, 1989; Ivry, Keele & Diener, 1988). In fact, CE lesion patients displayed larger heterogeneity in self-paced tapping and reduced capacities to synchronize their tapping to temporally regular sequences and tempo changes (Schwartze et al., 2016a). However, their interval discrimination was comparable to the one of healthy controls when tested in rhythmic contexts (Breska & Ivry, 2018, 2020, 2021). A reversed d pattern was observed in BG lesion patients, where interval discrimination deteriorated in rhythmic contexts (Breska & Ivry, 2018). This observation suggests a fine, yet fundamental dissociation between rhythm- and interval-based temporal predictions in BG and CE lesion patients (Breska & Ivry, 2018). BG patients’ timing performance is, however, quite heterogeneous: large variability is observed in interval timing tasks at both the second and millisecond range (Jones et al., 2008; Merchant et al., 2008), and they further show difficulties in coordinating and adjusting their tapping to tempo changes (Schwartze et al., 2011b). Supporting prior lesion studies, these observations suggest intact single interval-based timing in the BG patients (Breska & Ivry, 2018; Grube, Cooper, et al., 2010; Teki, Grube, Kumar, et al., 2011), but impaired processing of temporal regularity (i.e., the hierarchical temporal structure of a sensory sequence, the beat) in music (Grahn & Brett, 2009b) and speech (Kotz & Schmidt-Kassow, 2015).

What underlies these dysfunctions? Can we relate difficulties in producing and synchronizing with external rhythms to specific neural dynamics? In other words, can the analysis of neural oscillatory activity causally show in what way the cortico-subcortical timing network differentiates the processing of temporal regularities in perception and action?

We report results from two experiments involving healthy ageing controls (HC) and patients with focal lesions in either the CE or BG. Via a behavioral and an EEG experiment, we assessed sensory and sensorimotor capacities to encode, produce, and synchronize neural activity and overt behavioral responses with temporal regularities in auditory sequences.

Participants listened to isochronous auditory sequences while EEG was recorded. We assessed *if*, *when,* and *how* their neural activity reflected the temporal regularity of auditory sequences. We expected CE patients to show increased variability in the temporal encoding of tone onsets, while we hypothesized BG patients to have difficulties in generating temporal predictions. To test these hypotheses, we developed an extensive analysis pipeline designed to quantify the *encoding* and *tuning* of delta-band neural oscillatory activity towards the stimulation frequency (*Sf*).

First, we reproduced typical event-related potential (ERP) and inter-trial phase coherence (ITPC) analyses. The assessment of the N100 amplitude and latency along with the ITPC amplitude at the stimulation frequency (Sf; 1.5Hz) allowed testing if the participants’ neural activity responded in time to sound onsets, thus encoding their precise timing. In line with the evidence detailed above, we expected CE but not BG patients to show altered encoding of sound onsets: this would result in increased N100 latency variability and reduced ITPC. While ITPC and similar phase concentration measures are typically interpreted as a proxy of entrainment, it is generally hard to disentangle true entrainment from a sequela of evoked responses (Breska & Deouell, 2017b; Haegens & Zion Golumbic, 2018; Obleser et al., 2017; Zoefel et al., 2018). In fact, simulations showed that evoked responses drive high phase coherence (Breska & Deouell, 2017b; Obleser et al., 2017), ultimately weakening its functional link with temporal prediction. Thus, in a second analysis step, we introduced a novel approach to assess finer-grained measures of neural dynamics of temporal processing. First, we employed a time-resolved metric of ITPC (t-ITPC) to quantify the build-up of phase-coherence over the course of the auditory sequence. Such a metric was previously shown to be a good indicator of temporal prediction (Breska & Ivry, 2020) and auditory processing (Ten Oever et al., 2017). In fact, next to event-related t-ITPC peaks (which are induced by the phase rest of oscillations at tone onsets), the t-ITPC slope could indicate a mechanism of anticipation of next sound onsets. Thus, we expected BG but not CE patients to show a reduced slope of t-ITPC along the auditory sequence, potentially indicating altered temporal predictions. Next, we showcased a new method to assess the tuning of endogenous delta-band neural oscillatory activity towards the Sf. We quantified trial-level instantaneous frequency (IF), its dynamics of acceleration while tuning- and de-tuning to the Sf, its Stability (S), and Deviation (Dev) from the Sf. We expected BG and CE patients to show reduced S and greater Dev than HC, indicating their altered capacity to align neural dynamics to predictable onsets.

In the behavioral experiment, we expected the patient groups to display difficulties in producing stable finger-tapping and to synchronize with regular auditory sequences presented at three different tempi. We hypothesized that CE patients would have difficulties in the precise encoding of event onsets and synchronizing behavior, especially with faster tempi, while BG patients were expected to have more difficulties producing and synchronizing with slower rhythms (Nozaradan et al., 2017; Schwartze et al., 2011b). To test these hypotheses, we assessed individual tapping dynamics, quantifying tapping rate, acceleration, entropy, and stability of the performance over time.

## 2. Materials & Methods

### 2.1. Participants

Thirty-three participants took part in the study and signed written informed consent in accordance with the guidelines of the ethics committee of the University of Leipzig and the declaration of Helsinki. Participants comprised two patient groups and one age- and gender-matched control group: 11 patients with focal lesions in the basal ganglia (BG; mean age 50.9, range = 30-64 years; 5 males), 11 patients with focal lesions in the cerebellum (CE; mean age 52.6, range = 37-64 years; 5 males), and 11 healthy controls (HC; mean age 52.1, range = 28-63 years; 5 males). Patients’ demographics and lesion information are provided in Tab. 1, their anatomical MRI is provided in Fig. 1, and the results of their neuropsychological and cognitive assessment are provided in Suppl. Tab. 1.

**Figure 1.**
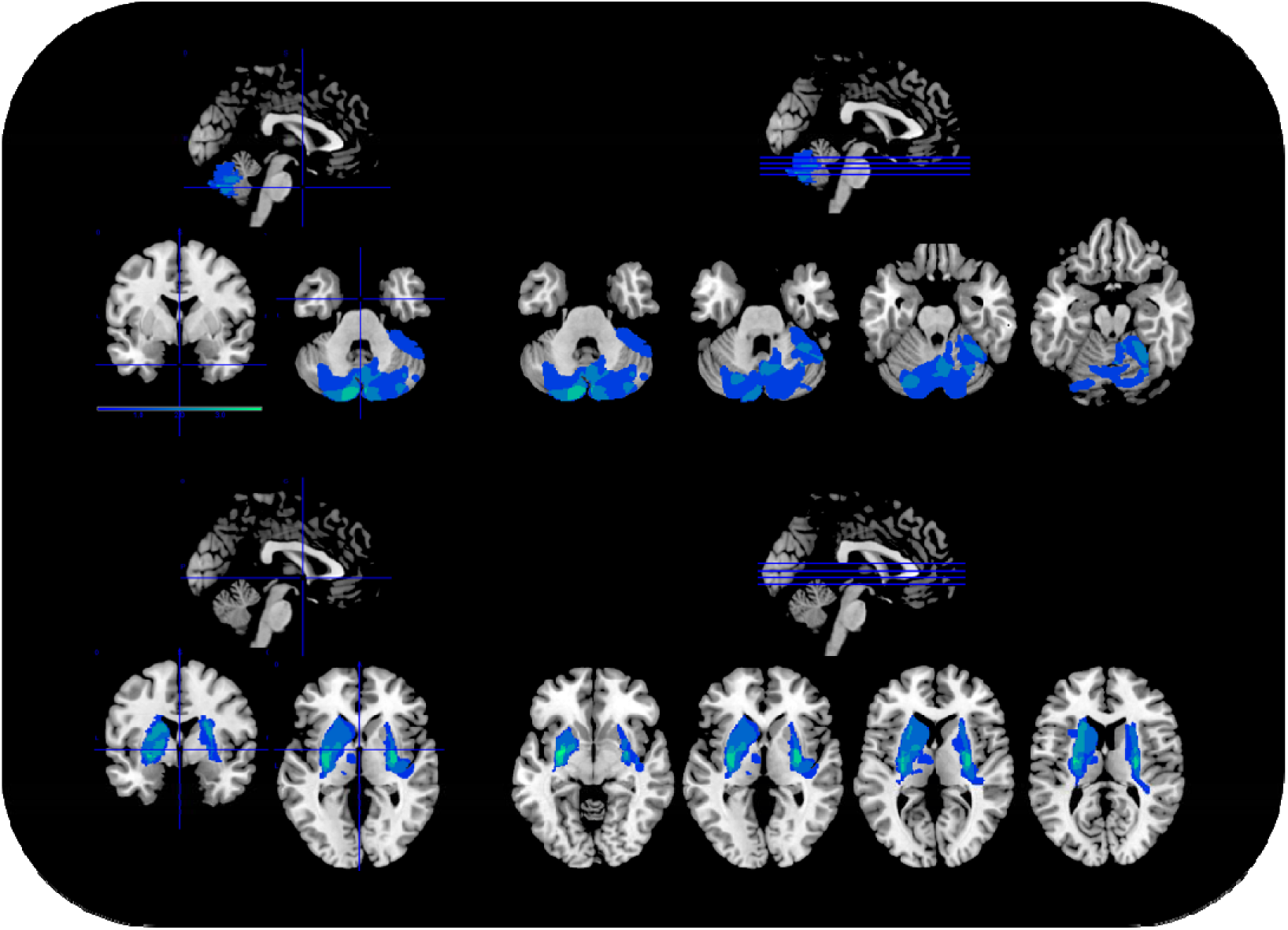
Brain lesion delineation and overlap. The figure provides the brain lesion delineation on a template anatomical MRI for cerebellar (CE; top) and basal ganglia (BG; bottom) patients. The figures were obtained by using MRIcron (http://www.mccauslandcenter.sc.edu/mricro/mricron/), an display the lesion overlap color-coded in shades of blue (blue = minimal overlap; light blue high overlap; values range from 0 t 5).

**Table 1.** Individual patient history. In order, from left to right, age, lesion location, secondary lesion location, lesion etiology, volume of the lesion in cc, and lesion coordinates for the center of mass (as provided by MRIcron). Top for cerebellar (CE) patients, and bottom for basal ganglia (BG) patients. Abbreviations: PICA = posterior cerebellar artery, ICH = intracerebral/-cerebellar hemorrhage, AVM = arteriovenous malformation, IC = internal capsule, EC = external capsule, MCA = Middle cerebral artery. The highlighted rows indicate patients for which both EEG and tapping data were used.

| ***CE*** | ***Age*** | ***Location*** | ***Additional locations*** | ***Etiology*** | ***Volume in cc*** | ***X*** | ***Y*** | ***Z*** |
| --- | --- | --- | --- | --- | --- | --- | --- | --- |
|  | 64 | posterior cerebellar lobule | none | PICA infarction | 7,1 | 89 | 37 | 26 |
|  | 30 | posterior cerebellar lobule | superior frontal gyrus, corpus callosum | ICH and AVM | 48 | 77 | 38 | 35 |
|  | 49 | posterior cerebellar lobule, tonsil, vermis | none | PICA infarction | 14,5 | 57 | 40 | 24 |
|  | 53 | anterior and medial cerebellar lobule, tonsil | none | PICA infarction | 7,9 | 62 | 41 | 24 |
|  | 39 | anterior and posterior cerebellar lobule | none | cyst extirpation | 2,4 | 88 | 52 | 15 |
|  | 62 | posterior cerebellar lobule | thalamus | PICA infarction | 9 | 111 | 69 | 37 |
|  | 62 | posterior cerebellar lobule | telencephalic white matter | PICA infarction | 3,1 | 57 | 58 | 55 |
|  | 59 | posterior cerebellar lobule, tonsil | none | PICA infarction | 3,6 | 110 | 38 | 30 |
|  | 63 | posterior cerebellar lobule | telencephalic white matter | PICA infarction | 5,7 | 104 | 61 | 46 |
|  | 37 | vermis, deep cerebellar nuclei, peduncle | none | tumor postoperatively | 8,8 | 84 | 49 | 37 |
|  | 42 | posterior cerebellar lobule | none | PICA infarction | 0,3 | 121 | 48 | 27 |
| **BG** |  |  |  |  |  |  |  |  |
|  | 41 | putamen, pallidum, caudate body, IC | corona radiata | MCA infarction | 23,4 | 56 | 113 | 76 |
|  | 59 | putamen, pallidum, caudate head and body, IC | corona radiata | ICH | 18,3 | 53 | 119 | 76 |
|  | 52 | putamen | none | ICH | 2,8 | 104 | 99 | 74 |
|  | 61 | claustrum, putamen, pallidum, caudate body, EC, IC | corona radiata, thalamus | ICH | 8,8 | 51 | 98 | 77 |
|  | 37 | putamen, caudate body, IC | none | MCA infarction | 0,8 | 55 | 97 | 88 |
|  | 50 | putamen, caudate body, IC | corona radiata, corpus callosum | MCA infarction | 6,7 | 54 | 110 | 81 |
|  | 60 | putamen | none | MCA infarction | 0,5 | 47 | 102 | 64 |
|  | 55 | putamen, pallidum, caudate body | none | MCA infraction | 3,0 | 102 | 101 | 77 |
|  | 51 | claustrum, putamen, EC | insula | ICH | 14,8 | 106 | 102 | 78 |
|  | 64 | putamen, pallidum, caudate body, IC | none | MCA infaction | 6,1 | 98 | 116 | 80 |
|  | 49 | putamen, caudate body | thalamus | MCA infarction | 6,3 | 93 | 103 | 82 |

The HC group was recruited via a database at the Max Planck Institute for Human Cognitive and Brain Sciences (Leipzig, Germany). All participants were right-handed and matched for years of education. None of the participants were professional musicians, and they reported no history of hearing or psychiatric disorders. All participants received monetary compensation for taking part in the study. Further information on the participants and lesions characteristics can be found in Nozaradan et al., (2017).

### 2.2. EEG experiment: design and procedure

Participants listened to 96 sequences comprising 13-to-16 tones in a recording session of approximately 25min. Each sequence contained frequent standard tones (STD; F0 = 400Hz, duration = 50ms, rise and fall times = 10ms, amplitude = 70dB SPL) and one or two amplitude-attenuated deviant tones (DEV; amplitude 66dB). The inter-onset-interval between successive tones was 650ms, resulting in a stimulation frequency (*Sf*) of 1.54Hz, and a total sequence duration of 8.45-10.4s (13 to 16 tones * 650ms; Fig. 2A).

**Figure 2.**
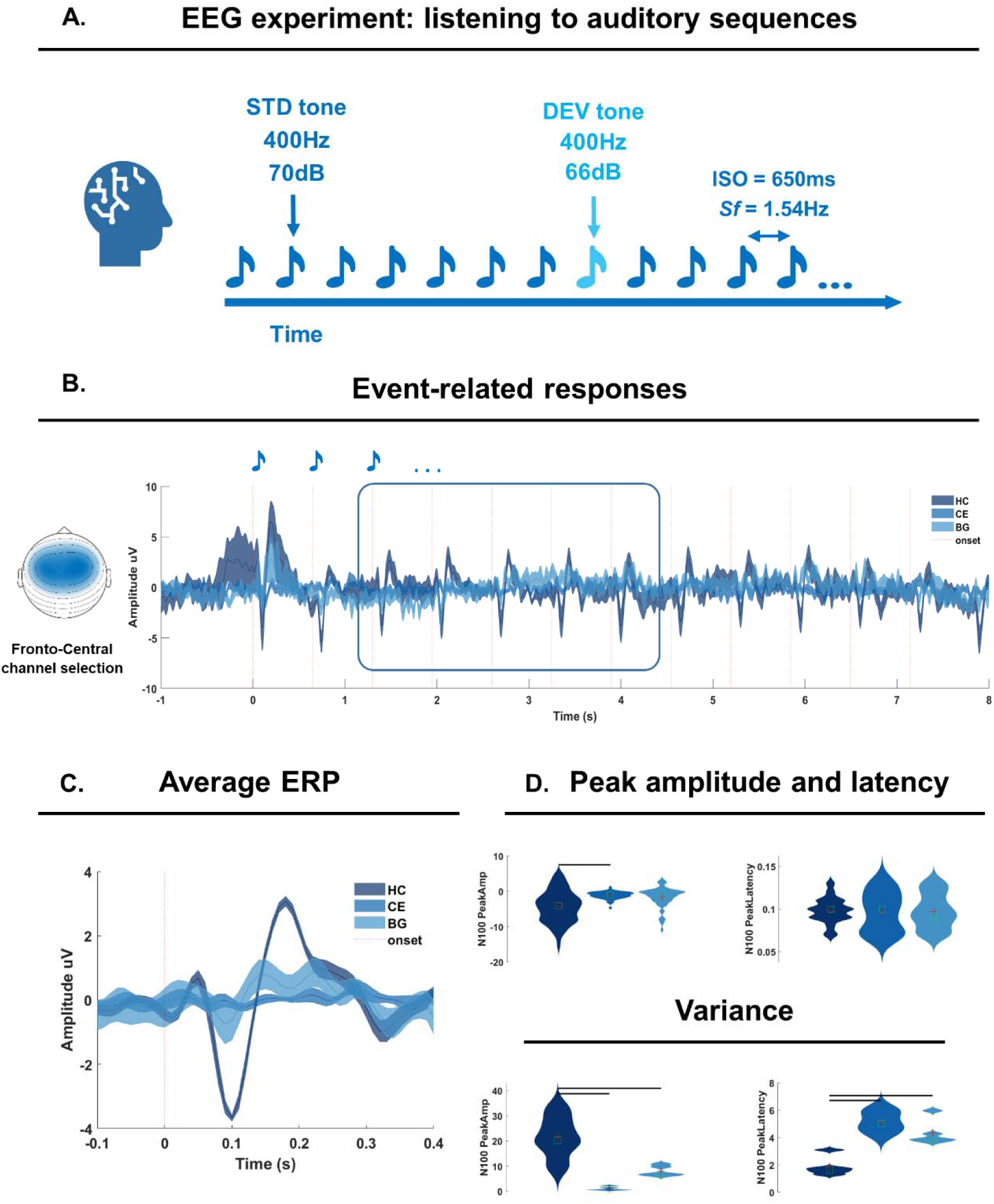
EEG experiment and ERP analyses. A: experimental design: participants listened to 96 isochronous auditory sequences presented at a fixed Stimulation frequency (Sf) of 1.54 Hz (inter-stimulus onset (ISO) of 650ms). The sequences included frequent standard (STD; pitch 400Hz, loudness 70dB) and infrequent amplitude deviant (DEV) tone (amplitude 66dB). B: sequence-level ERPs as calculated in a fronto-central channel cluster (left) and color coded per group (dark blue for HC; light blue for BG; medium blue for CE). C: tone-level ERP, obtained by averaging across sound repetitions, and among tones in the 3^rd^ to 7^th^ position (as highlighted by the rectangle in 2B). D. Group-level peak amplitude (top left) and latency (top right). Below, group-level variance in peak amplitude and latency. Horizontal lines report a significant group effect and significant pairwise comparisons (see Tab 2,3,4).

Participants were seated in a dimly lit soundproof chamber facing a computer screen. Every trial started with a fixation cross (500ms), followed by the presentation of an auditory sequence. The cross displayed on the screen served to prevent excessive eye movements during the presentation of the tone sequences. After each sequence, there was an inter-trial interval of 2000ms. A session was divided into two blocks of approximately 10 minutes each, with a short pause in between. At the end of the experiment, participants indicated how many softer tones they heard.

#### 2.2.1. EEG recording

The EEG was recorded by means of 59 Ag/AgCl scalp electrodes positioned according to the International 10-10 system with the ground placed on the sternum. Four additional vertical and horizontal electrodes monitored eye movements and were positioned on the outer canthus of each eye, and on the inferior and superior areas of the left orbit. The signals were amplified, low-pass filtered at 512Hz and digitized using a sampling rate of 1024Hz (64-channel high-speed amplifier, Biosemi, the Netherlands). Electrode impedances were kept below 5kΩ and the left mastoid served as online reference. Data were referenced to an average reference offline.

#### 2.2.2. Data Analysis

##### EEG Preprocessing

EEG data were analyzed in MATLAB with a combination of custom scripts and functions and the FieldTrip toolbox (Oostenveld et al., 2011). Data were band-pass filtered with a 4th order Butterworth filter in the frequency range of 0.1-50Hz (*ft_preprocessing*). Eye-blinks and other artifacts were identified using independent component analysis. This semi-automated routine is composed of two steps: in the first iteration, ‘*fastICA’* (implemented in FieldTrip) was applied to decompose the original EEG signal into independent components (N= number of EEG channels - 1). Components with a strong correlation (>.4) with the EOG time-courses were automatically identified and removed with ‘*ft_rejectcomponent*’ before reconstructing the EEG time-course. In a second step, ‘*fastICA’* was used again, now with a dimensionality reduction to 20 components. These components were visually inspected via ‘ft_*rejectvisual’* and marked as ‘outliers’ if their max values and z-scores were far from the distribution of other components. The 20 components were further visually inspected by plotting their topographies and time-series, and a second selection of ‘outliers’ was made. Taking into consideration the two visual inspections, we made a final decision on which components to remove. On average, 2 components were removed (‘*ft_rejectcomponent*’) before reconstructing the EEG time-series. In the next preprocessing step, artifact subspace reconstruction was performed as implemented in the ‘*pop_clean_rawdata*’ function in EEGlab, and with the ‘BurstCriterion’ parameter set to 20 (as recommended in the online EEGlab tutorials; all other parameters were set to ‘off’). To further ensure the removal of potentially noisy channels and time-points, we implemented an automatic channel rejection and an artifact suppression procedure. To this end, the median variance across channels was calculated (excluding EOG channels), and channels exceeding 2.5*median variance were defined as ‘outliers’ and removed. Similarly, the artifact suppression procedure (see Criscuolo et al., 2023) interpolated noisy (>4*absolute median) time-windows on a channel-by-channel basis. Lastly, data were low pass filtered at 40Hz via ‘*ft_preprocessing*’, segmented to each auditory sequence (starting 4s before the first tone onset and ending 4s after the last tone onset), and downsampled to 250Hz. All the following analyses focused on a region of interest. As in prior work (Criscuolo et al., 2023), a fronto-central channel (FC) cluster was used, encompassing the sensor-level correspondents of prefrontal, pre-, para-, and post-central regions highlighted in (Fujioka et al., 2012; Fujioka et al., 2015) and further highlighted in similar EEG work on rhythm processing (Nozaradan et al., 2012). The cluster included 14 channels: ‘AFz’, ‘AF3’, ‘AF4’, ‘F3’, ‘F4’, ‘F5’, ‘F6’, ‘FCz’, ‘FC3’, ‘FC4’, ‘FC5’, ‘FC6’, ‘C3’, ‘C4’.

##### ERP analyses

To perform ERP peak analyses, preprocessed data was low-pass filtered at 30Hz via ‘*ft_preprocessing*’ and averaged across FC channels. ERP time-series are provided in Fig. 2B, color-coded for each group. Next, these time-series were segmented to each tone onset in the auditory sequences (average over trials provided in Fig. 2C). To perform peak analyses, we set out to identify, at the single participant- and trial-level, the N100 peak amplitude and its latency for each tone onset. For doing so, we created a window of interest in the time interval between .07s and .13s relative to tone onsets, and identified the most negative amplitude value (N100 peak amplitude) and its latency (N100 peak latency). Finally, we calculated, at the single-participant level, the variance in N100 amplitude and latency. Peak amplitude, latency, and variance are provided in Fig. 2D, pooling single-participant data per group. Analyses focused on the 3^rd^ to the 7^th^ tone: the first two sound onsets were excluded as they are known to trigger much stronger ERPs; the later tones are excluded because they occur in proximity of the DEV (8^th^ position onward).

###### Mixed Effect Models on ERP data

We assessed group differences in the N100 peak amplitude and latency via Mixed effect models. The model included ‘Group’ as a fixed factor and a random intercept per participant. When the fixed effect was significant, we performed post-hoc pairwise group comparisons via permutation testing. Pair-wise comparisons were thus performed by randomly permuting data points belonging to one or the other group, with a total of 1000 permutations. This iterative procedure would ultimately assess the p-value from the original groups against the p obtained from permutations. Model information and results are reported in Tab 2-3.

###### Statistical comparisons on the variance

We assessed the variability in the N100 peak amplitude and latency across groups. For these two analyses, we first performed Levene’s test to assess the homogeneity of variance and later employed ANOVA. With a significant group effect, we later performed post-hoc pairwise comparisons via simple t-tests and implementing the Tukey-Kramer correction. Analyses details and results are provided in Tab 4-5.

##### Inter-Trial Phase coherence

Inter-trial phase coherence (ITPC) analyses were conducted to test whether healthy participants and patients encoded temporal regularities in the auditory sequences. To estimate ITPC, we first performed Fast-Fourier transform (FFT). FFT analyses were performed at the single-participant, -channel and -trial level on 8s-long segments starting from the onset of the first tone in the auditory sequence and including a total of 12 tones. The resulting frequency resolution was .125Hz (1/8s = .125Hz). As in prior work (Criscuolo et al., 2023), and for the ERP analyses described above, a fronto-central channel (FC) cluster was used, including the following 14 channels: ‘AFz’, ‘AF3’, ‘AF4’, ‘F3’, ‘F4’, ‘F5’, ‘F6’, ‘FCz’, ‘FC3’, ‘FC4’, ‘FC5’, ‘FC6’, ‘C3’, ‘C4’. Data from this FC cluster were not averaged at this stage. Next, the complex part of the Fourier spectrum was used to calculate ITPC (Fig. 3A left). ITPC was obtained by dividing the Fourier coefficients by their absolute values (thus, normalizing the values to be on the unit circle), averaging, and finally taking the absolute value of the complex mean (for further documentation see https://www.fieldtriptoolbox.org/faq/itc/). For illustration purposes, the ITPC plot in Fig. 3A is restricted to 1-4Hz.

**Figure 3.**
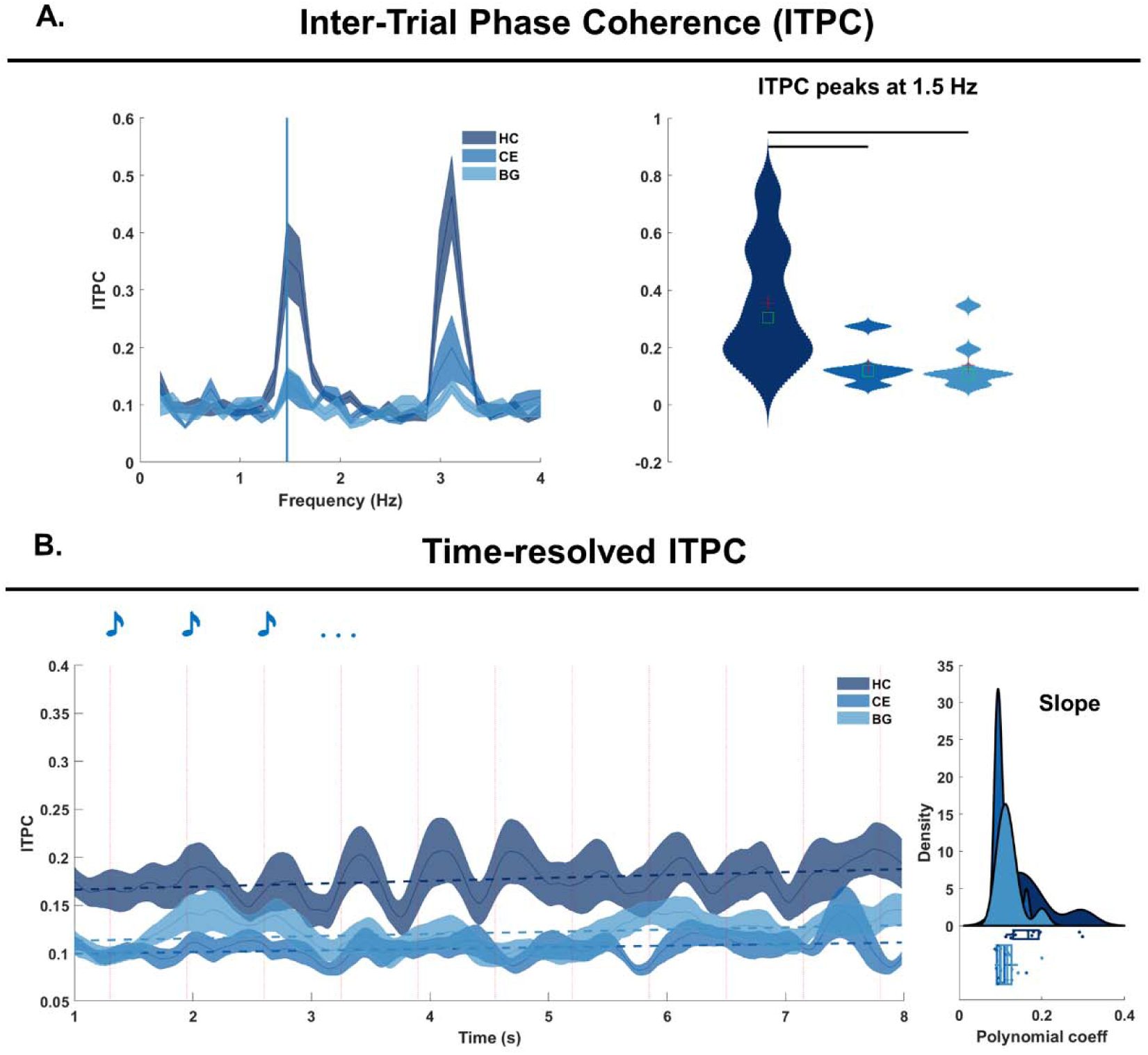
Inter-Trial Phase Coherence (ITPC) analyses. A: Inter-Trial Phase Coherence (ITPC) analyses. The ITPC plot features frequency in Hz on the x-axis and the ITPC values o the y-axis. Healthy controls (HC), cerebellar (CE) and basal ganglia (BG) patients’ data are provided in shades of blue (from dark to light blue). The vertical blue line signals the ITPC peak at 1.5Hz. These peaks were extracted and plotted on the right, via means of violin plots. Here, statistical comparisons assessed group differences in ITPC values at the stimulation frequency (1.5Hz). The black horizontal lines indicate significant differences between the groups, after correction for multiple comparisons. B: time-resolved ITPC was computed from the complex spectra of continuous wavelet transformed data. The left plot features the time-course of t-ITPC in the time-range between 1-8s (x-axis) and the ITPC value on the y-axis. On top, the dashed lines represent the slope of each time-course. Each participant’s slope is then plotted in the distribution on the right side: polynomial coefficient on the x-axis and density on the y-axis.

As the stimulation frequency of the auditory sequences was 1.5Hz, we restricted ITPC analyses to 1.5Hz only. We did not conduct any statistical analyses on any other frequencies (e.g., (sub)harmonics of the stimulation frequency) because we were specifically interested in assessing the neurophysiological encoding of the auditory rhythm at the stimulation frequency as previously reported in Criscuolo et al., 2023a; Criscuolo et al., 2023b. Thus, single-participant data were pooled per group after averaging across channels and visualized as violin plots in Fig. 3A (right).

###### Statistical comparisons

Group differences were statistically assessed by means of a 1-way ANOVA with a group factor (*anova1* built-in in MATLAB), followed by post-hoc simple tests corrected for multiple comparisons via Tukey-Davis correction (*multcompare* built-in in MATLAB) in case of a significant (*p* <.05) main effect (Tab. 6). Simple effects with a *p-*value below an alpha-corrected .05 were considered statistically significant. The choice of the parametric test was informed by Levene’s test for homogeneity of variance (Tab. 6).

##### Time-resolved ITPC

A time-resolved metric of ITPC (t-ITPC) was estimated to quantify the build-up of phase-coherence over the course of an auditory sequence. For doing so, we first employed time-frequency transform (TF data), then calculated the t-ITPC using the complex spectra of TF data (same procedure as in the ITPC above), and finally calculated the slope of the t-ITPC at the single-participant level. More details are provided in the respective paragraphs below.

###### Time-frequency transform

After preprocessing, single-trial EEG data underwent time-frequency transformation (‘*ft_freqanalysis’*) by means of a wavelet-transform (Cohen, 2014). The bandwidth of interest was centered around the stimulation frequency (+/- 1Hz, i.e., .54 - 2.54Hz, thus obtaining a 1.54Hz center frequency), using a frequency resolution of .2Hz. The number of fitted cycles was set to 3. The single-trial approach results in ‘induced’ (as compared to ‘evoked’) responses. The output was a complex spectrum; no averaging over channels, trials, or participants was performed at this stage.

###### Slope of t-ITPC

As for the ITPC above, t-ITPC was obtained by dividing the complex coefficients of TF-data by their absolute values (thus, normalizing the values to be on the unit circle), averaging, and finally taking the absolute value of the complex mean. Next, we calculated the slope of each t-ITPC time-series by fitting a first-order polynomial (‘*polyfit*’ function in MATLAB; *p*) and then deriving a first-order approximation (*p(1)*Time+p(2)*). The calculation of the slope was performed on a 5s-long interval starting from the 3^rd^ tone onset.

##### Statistical comparisons

Before assessing group differences in the t-ITPC, we performed Levene’s test for homogeneity of variance. As the *p* returned significant, we implemented non-parametric group comparisons via a Kruskal-Wallis test. With a significant group effect, we later assessed pairwise group comparisons via permutation testing and with a total of 1000 permutations. This iterative procedure would ultimately assess the p value from the original groups against the p obtained from permutations. Analysis steps and results are reported in Tab 7.

##### Analyses of oscillatory dynamics

We aimed at assessing the tuning of endogenous delta-band neural oscillatory activity towards the stimulation frequency (*Sf*). A self-sustained endogenous oscillator capable of entrainment should adjust its frequency and phase so to align to external temporal regularities, or rhythms.

Such an active process is displayed in tuning dynamics during which the endogenous oscillator accelerates and decelerates to align its period to an external rhythm. Thus, we calculated the instantaneous frequency (IF) and quantified acceleration dynamics. While processing the auditory rhythm, oscillations may be fluctuating around the *Sf* with various magnitude and deviating from the *Sf*. So, we calculated metrics of stability and deviation. Details for each of the metrics are provided in the respective sections below.

###### Instantaneous frequency

After preprocessing, single-trial EEG data were bandpass-filtered with a 3rd order Butterworth filter centered around the stimulation frequency (plus and minus 1Hz, .54 - 2.54Hz, obtaining a 1.54Hz center frequency; *ft_preprocessing*) and Hilbert-transformed to extract the analytic signal. Next, the instantaneous frequency (IF) at each time point (t), and for each channel, trial, and participant, can be calculated with the following formula:

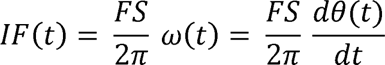

Where *ω*(t) is the derivative of the unwrapped phase (*θ*) at each time point (t), given the time-steps (*dt*) and FS is the sampling frequency. IF can also be obtained in MATLAB using the ‘*instfreq*’ function.

###### Acceleration

Once calculated the single-participant, -trial, and channel-level IF, acceleration (Acc) was calculated as the first derivative of IF. Thus, we employed the following formula:

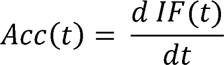

###### Stability

Once the single-participant, -trial, and channel-level Acc was obtained, Stability (S) was calculated as the inverse of the sum of absolute changes in Acceleration. Thus, we employed the following formula:

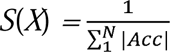

###### Deviation

Once the single-participant, -trial, and channel-level IF was obtained, we quantified the standard deviation (D) from the stimulation frequency. D was calculated as the square-root of the mean squared difference between the IF and the stimulation frequency (*Sf*):

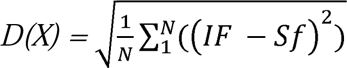

###### Latency

For each participant, trial, and channel, we estimated how long it took for the IF to tune to the *Sf* (1.5Hz). The search was performed within the first 8s of listening (0-8s) of each trial, and by using a narrow frequency criterion (*Sf ±* .2Hz).

###### Statistical comparisons

Group differences were statistically assessed for each of the above-mentioned metrics by means of a 1-way ANOVA with a group factor (*anova1* built-in into MATLAB), followed by post-hoc simple tests corrected for multiple comparisons (Tukey-Davis correction) (*multcompare* built-in into MATLAB) in case of a significant (*p* <.05) main effect. Simple effects with a *p-*value below an alpha-corrected .05 were considered statistically significant and are depicted as horizontal black lines on top of Fig. 4B. The ANOVA tables and simple tests for significant effects are provided in Tab. 7.

**Figure 4.**
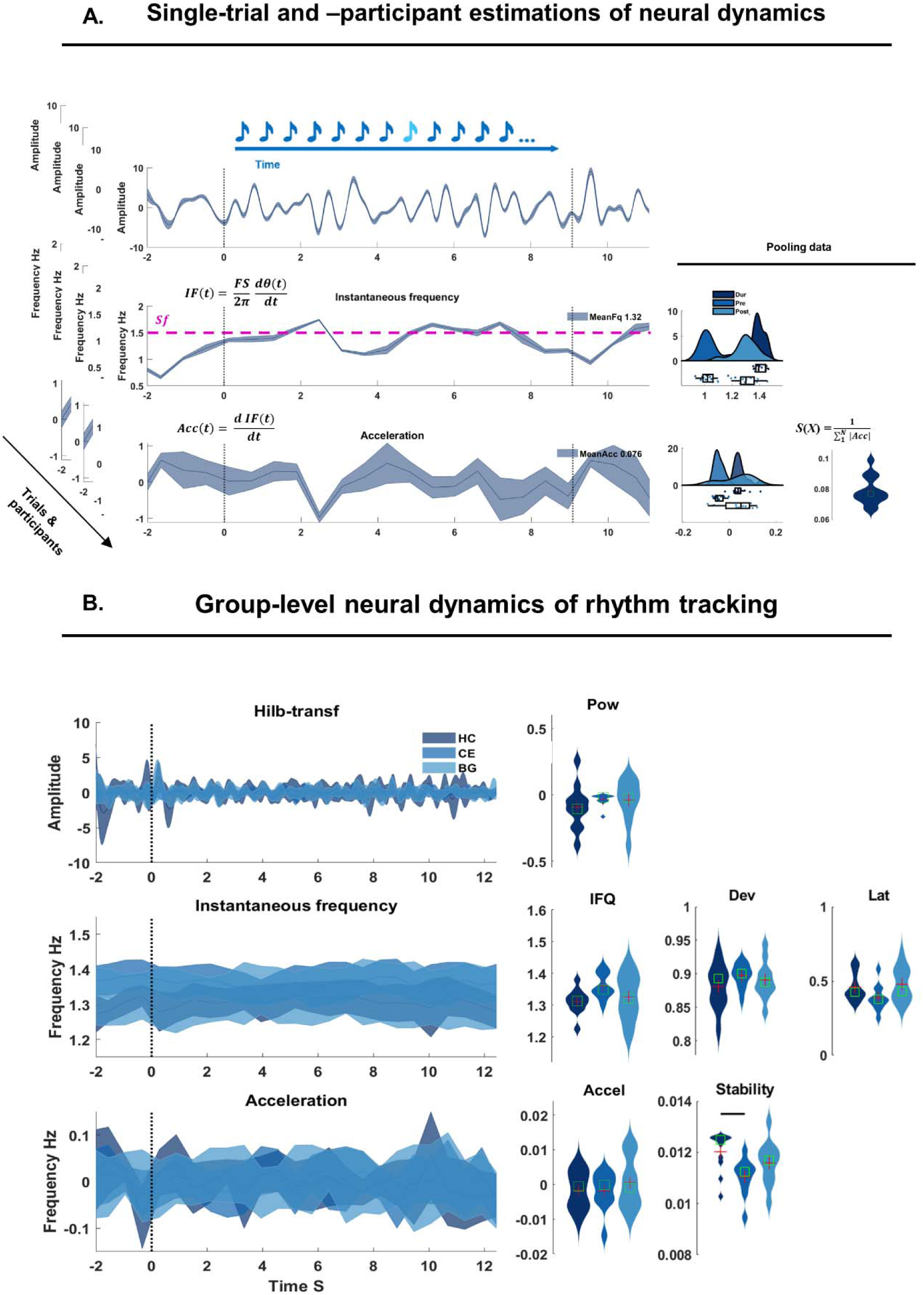
Analyses on neural dynamics of rhythm processing. A: After obtaining the analytical signal of delta-band activity (top), we estimated, at the single-participant and -trial levels, the instantaneous frequency (IF) and Acceleration (Acc; time-courses on the left side). The pink dashed line on top of the IF indicates the stimulation frequency (Sf). The distribution plots on the right side are obtained by pooling data points over channels and averaging over time. IF and Acc were also used to calculate metrics of Deviation (D) and Stability (S). The violin plots on the right side show the resultant D and S over time, and pooling over channels. B: mirroring the layout of A, the group-level data (color-coded in shades of blue) reporting on the left the time-courses of neural oscillations in the delta frequency band (top), the estimated IF (middle) and Acc (bottom). On the right side of the IF time-courses, violin plots of the IF (pooling over participants, but averaging over trials and across the FC cluster of interest). Next to it, the single-participant Dev and Latency. Next to the time-courses of the Acc, violin plots of the Acc (pooling participants and averaging across trials and the FC cluster of interest). On its right, the estimated S values. The thick black horizontal lines on top of the violin plots indicate significant group differences.

### 2.3. Tapping experiment

The same participants also took part in a tapping experiment. Participants listened to the same auditory stimuli as the ones employed in an earlier EEG study (Nozaradan et al., 2017). Sequences were composed of 12 intervals of 200ms and comprised pure tones (1000Hz) with 10ms rise and fall times and silent intervals (Fig. 5A). Such short inter-onset intervals (IOI) define a basic stimulation frequency (*Sf*) of 5Hz. The sequences were further presented at two more tempi: double (resulting in a 10Hz *Sf*), and quadruple (resulting in a 20Hz *Sf*) base tempo. Each sequence (12 intervals) was looped continuously for 33s, resulting in a total duration of ∼5.5min. Participants were asked to tap with the index finger in time with the perceived beat in an auditory stream and were instructed to start tapping as soon as they perceived the beat. Tapping was performed on a response pad while the forearm and elbow were fixed on an armrest cushion to avoid excessive movements. Taps did not trigger any sound, and the latency of each finger tap (i.e., the time of contact of the finger onto the pad) was registered with millisecond accuracy and recorded in Presentation (NBS, Berkeley, USA).

**Figure 5.**
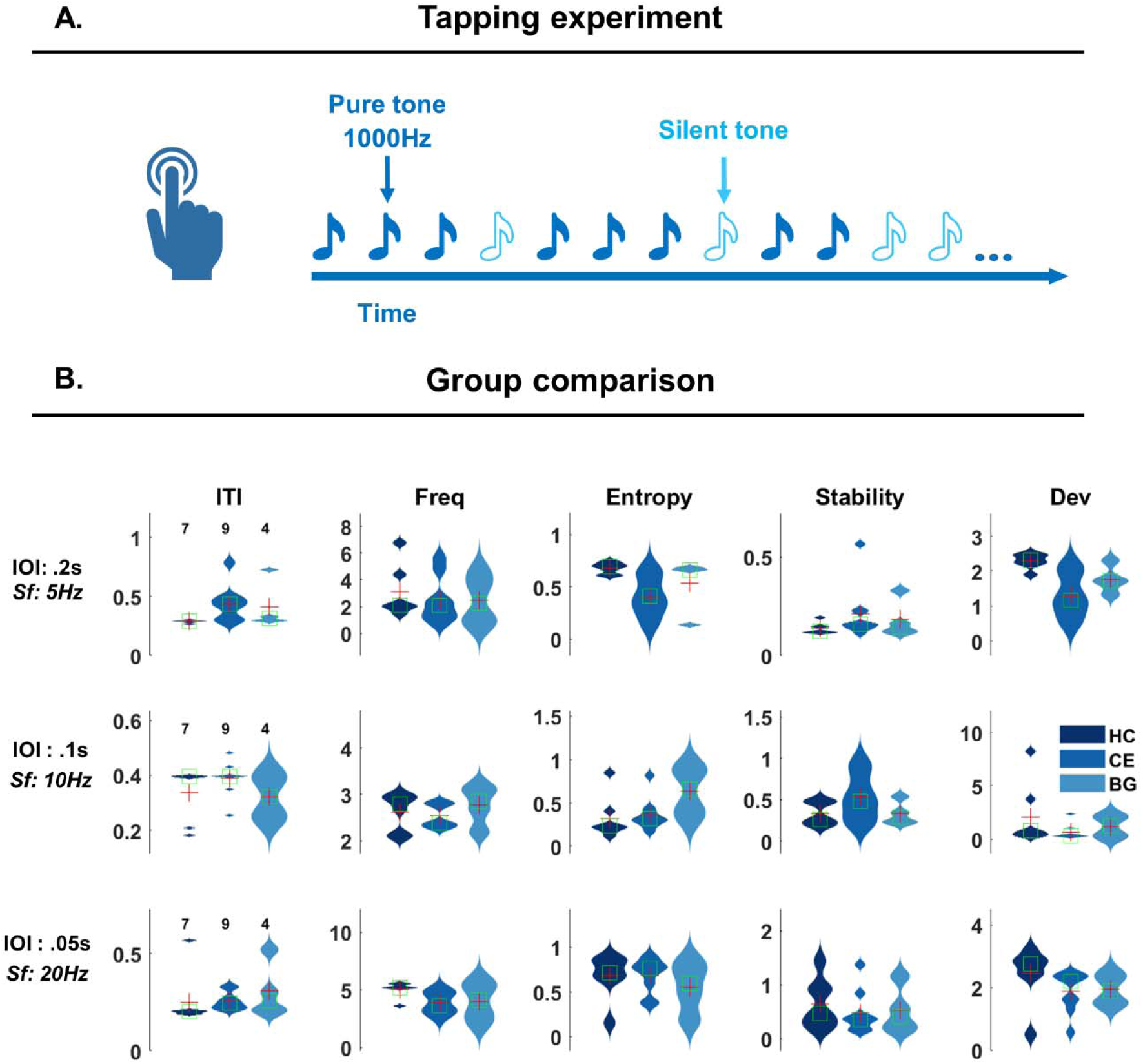
Tapping experiment and group data. A: experimental design: participants listened to auditory sequences at three different speeds and were asked to synchronize their tapping to the perceived beat, while keeping the tapping at a comfortable rate. These sequences included pure tones at 1000Hz and silent tones. The inter-onset-intervals (IOIs) varied in three steps: the basic tempo had IOI of .2s,and the other tempos had IOIs of.1s and.05s respectively, resulting in stimulation frequencies (Sf) of 5, 10 and 20 Hz. B: group data for inter-tap-intervals (ITI), Frequency, Shannon Entropy, Stability and Deviation, color coded per group in shades of blue (darker blue from healthy controls and lighter blues for CE and BG patients). On top, data for the 5Hz tempo; middle row for the 10Hz tempo; bottom row for the 20Hz tempo.

The speeding-up of auditory sequences served to increase task difficulty: to synchronize tapping to the stimulus material, participants had to *detect* the temporal regularities in the sequences, and coordinate motor behavior to *produce* and *synchronize* their tapping to a sub-harmonic of the *Sf*.

However, not all participants could complete the tapping task. We therefore present data from participants with at least 40 taps. Given the reduced number of data points, participants and the different number of participants per group, these data will not be used for statistical comparisons but only for a qualitative synthesis. The participants included in the analyses are highlighted in Tab. 1.

#### 2.3.1. Tapping data analysis

Based on tapping time data, we obtained the inter-tap-interval (ITI) as the difference between successive tap onsets. Next, we calculated the tapping frequency (Freq) as the inverse of the ITI (1/ITI), the Stability (with the same formulas as described above), Shannon Entropy, and Deviation. The latter was calculated with the same formula as mentioned above, as a standard deviation from 2.5Hz (a subharmonic of the stimulation frequency).

Individual data were pooled for each group (Fig. 5B). No statistical comparisons were performed due to low statistical power.

### 2.4. Data and code Availability

The analysis code in use will be stored in an open repository.

## 3. Results

In this study, we investigated the causal role of CE and BG lesions on temporal processing. Hence, we report results from two experiments: first, participants listened to isochronous auditory sequences while EEG was recorded continuously (Fig. 2A). Next, the same individuals performed a tapping task, in which they synchronized their tapping to regular auditory sequences presented at three different tempi (Fig. 5A).

### 3.1. EEG experiment

In this experiment, we adopted two complementary methodological approaches.

First, we reproduced typical event-related potential (ERP) and inter-trial phase coherence (ITPC) analyses to test *if* and *how* participants’ neural activity encoded the temporal regularity of sound onsets. In the second analysis step, we employed a time-resolved metric of ITPC (t-ITPC) to quantify the build-up of phase-coherence over the course of the auditory sequence. the t-ITPC slope could indicate a mechanism of phase-alignment towards next sound onsets (Breska & Ivry, 2020; Ten Oever et al., 2017). Next, we introduced a new method to assess the tuning of endogenous delta-band neural oscillatory activity towards the *Sf*. We quantified trial-level instantaneous frequency (IF), its dynamics of acceleration while tuning- and de-tuning to the Sf, its Stability (S), and Deviation (Dev) from the Sf.

Details for each methodological approach can be found in the respective methods section above. Results for each method are discussed below.

#### 3.1.1. Event-related potential (ERP) analyses

After computing the sequence-level (Fig. 2B) and tone-level ERPs (Fig. 2C), we obtained single-participant and -trial level N100 peak amplitude, latency (Fig. 2D, top) and later estimated their variance (Fig. 2D, bottom). Participant-level N100 peak amplitude and latency separately entered a Mixed Effect Model with a Group Factor and a random intercept per participant. The first model reported a significant group effect for N100 peak amplitude (F(2,165) = 2.33, *p* = .02; Tab 2). Post-hoc pairwise comparisons showed that CE had a lower N100 amplitude peak than HC (Obs Diff = -3.23, *p* = .01, Eff size = -1.33). There were no significant differences between HC and BG, nor between CE and BG (Tab. 2).

**Table 2.**
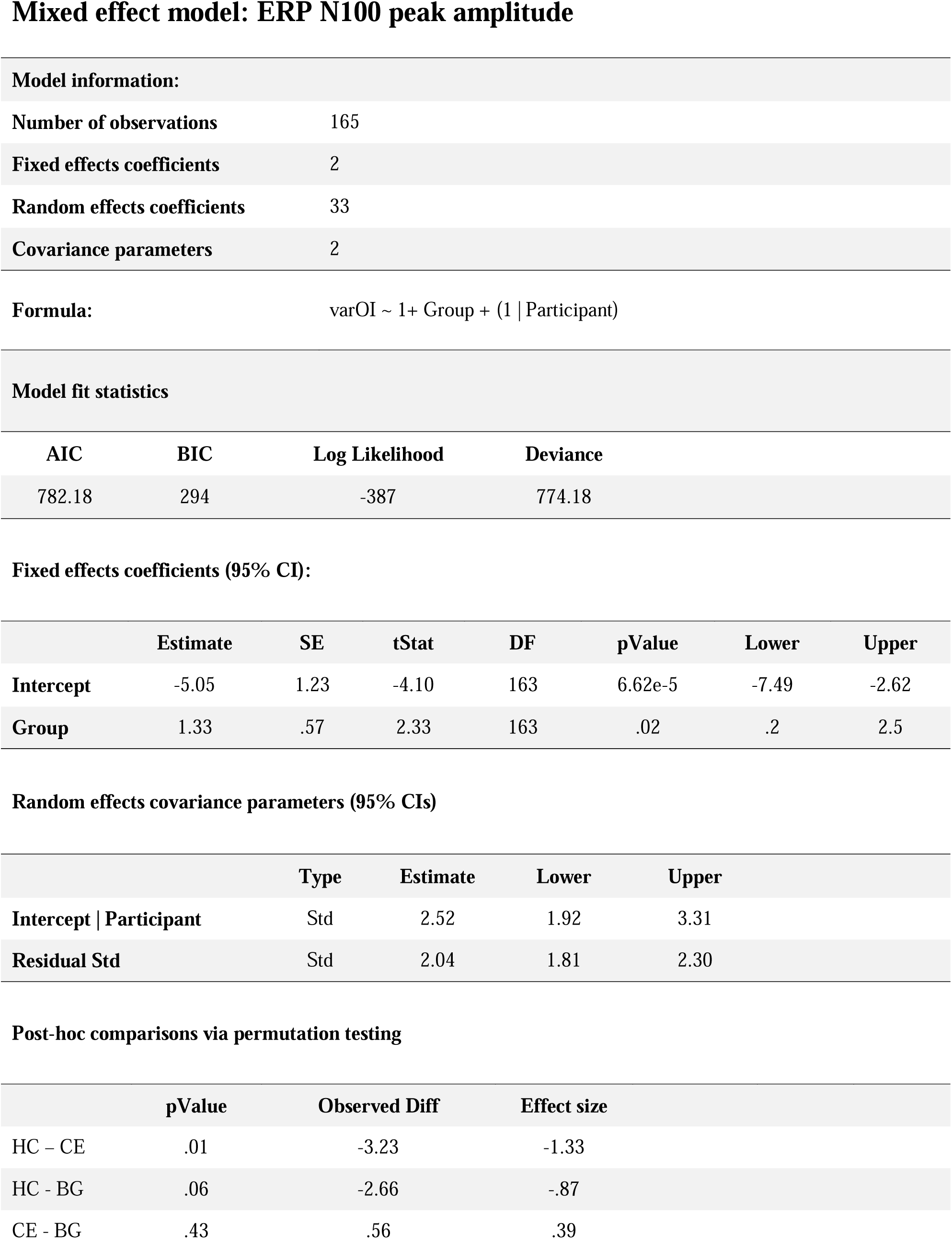
Mixed effect model on ERP N100 Peak Amplitude. The table reports model information: number of observations, fixed effect coefficients, random effect coefficients, covariance parameters. Then, the formula used to fit the model and model fit statistics: AIC, BIC values, Log Likelihood and Deviance. Further below, the fixed effect coefficients in a 95% confidence interval (CI): estimate, standard error, t-stat, degrees of freedom (DF), p value, lower and upper bound. Right below, random effects covariance parameters: estimate, lower and upper bound. At the bottom, the results of post-hoc comparisons performed via permutation testing: p-value, observed difference and effect size.

Model fit statistics
| AIC | BIC | Log Likelihood | Deviance |
| --- | --- | --- | --- |
| 782.18 | 294 | -387 | 774.18 |

Fixed effects coefficients (95% CI):
|  | Estimate | SE | tStat | DF | pValue | Lower | Upper |
| --- | --- | --- | --- | --- | --- | --- | --- |
| Intercept | -5.05 | 1.23 | -4.10 | 163 | 6.62e-5 | -7.49 | -2.62 |
| Group | 1.33 | .57 | 2.33 | 163 | .02 | .2 | 2.5 |

Random effects covariance parameters (95% CIs)
|  | Type | Estimate | Lower | Upper |
| --- | --- | --- | --- | --- |
| Intercept \| Participant | Std | 2.52 | 1.92 | 3.31 |
| Residual Std | Std | 2.04 | 1.81 | 2.30 |

Table 2 – Mixed effect model on ERP N100 Peak Amplitude
|  | pValue | Observed Diff | Effect size |
| --- | --- | --- | --- |
| HC – CE | .01 | -3.23 | -1.33 |
| HC - BG | .06 | -2.66 | -.87 |
| CE - BG | .43 | .56 | .39 |

The second model reported no significant group effect for N100 peak amplitude latency (F(2,165) = -.16, *p* = .88; Suppl. Tab 1). Next, we assessed the variability in the N100 peak amplitude. As the Levene’s test for homogeneity of variance was not significant (Tab. 3, top), we employed one-way ANOVA with a group factor. The ANOVA reported a significant group effect (F(2,30) = 21.26, *p* < .001; Tab. 3, middle) and pairwise post-hoc comparison revealed that HC had higher variance than either CE or BG, but there was no difference between CE and BG (Tab. 3, bottom). Finally, we assessed the variability in N100 peak amplitude latency. As the Levene’s test for homogeneity of variance was not significant (Tab. 4, top), we employed one-way ANOVA with a group factor. The ANOVA reported a significant group effect (F(2,30) = 22.14, *p* < .001; Tab. 4, middle) and pairwise post-hoc comparison revealed that both CE and BG had higher variance than HC but there was no difference between CE and BG (Tab. 4, bottom).

**Table 3.** Group comparison for the N100 peak amplitude variance. In order, the table reports results from the Levene’s test for homogeneity of variance: F-statistics, degrees of freedom (DF) and p value. In the middle, the results from the ANOVA: sum of squares (SS), degrees of freedom (DF), mean squared error (MS), F-statistics and p-value. The error term reports the within-group variation. At the bottom, the results of the pairwise group comparison, with Tukey-Kramer correction. In order, the p-value, observed difference, lower and upper limit.

ERP N100 peak amplitude variance
| Levene's test for homogeneity of variance |  |  |  |
| --- | --- | --- | --- |
|  | F stat | DF | pValue |
| **Group** | 3.73 | 2, 30 | .06 |

| Group Comparison via parametric test: ANOVA |  |  |  |  |  |
| --- | --- | --- | --- | --- | --- |
|  | SS | DF | MS | F stat | pValue |
| **Group** | 1090 | 2 | 545 | 21.26 | <.001 |
| **Error** | 308 | 30 | 26 |  |  |

Table 3 – Group comparison for the N100 peak amplitude variance.
| Post-hoc comparisons with Tukey-Kramer correction |  |  |  |  |
| --- | --- | --- | --- | --- |
|  | pValue | Observed Diff | Lower Limit | Upper Limit |
| HC – CE | 9.2e-5 | 20.54 | 12 | 29 |
| HC - BG | .003 | 13.53 | 4.99 | 22 |
| CE - BG | .11 | -7 | -15.6 | 1.54 |

**Table 4.**
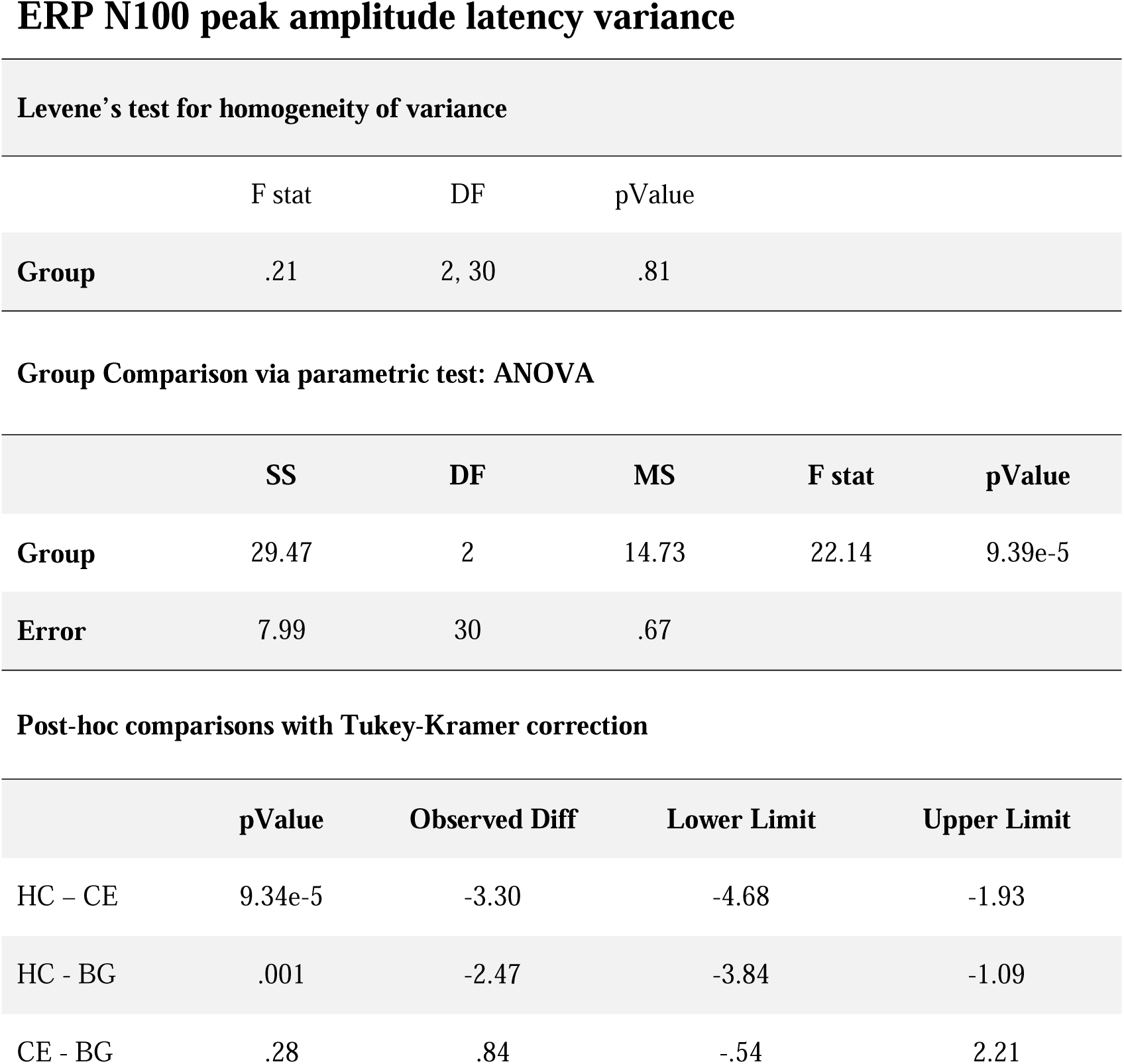
Group comparison for the N100 peak amplitude latency variance. In order, the table reports results from the Levene’s test for homogeneity of variance: F-statistics, degrees of freedom (DF) and p value. In the middle, the results from the ANOVA: sum of squares (SS), degrees of freedom (DF), mean squared error (MS), F-statistics and p-value. The error term reports the within-group variation. At the bottom, the results of the pairwise group comparison, with Tukey-Kramer correction. In order, the p-value, observed difference, lower and upper limit.

ERP N100 peak amplitude latency variance
| Levene's test for homogeneity of variance |  |  |  |
| --- | --- | --- | --- |
|  | F stat | DF | pValue |
| **Group** | .21 | 2, 30 | .81 |

| Group Comparison via parametric test: ANOVA |  |  |  |  |  |
| --- | --- | --- | --- | --- | --- |
|  | SS | DF | MS | F stat | pValue |
| **Group** | 29.47 | 2 | 14.73 | 22.14 | 9.39e-5 |
| **Error** | 7.99 | 30 | .67 |  |  |

Table 4 - Group comparison for the N100 peak amplitude latency variance.
| Post-hoc comparisons with Tukey-Kramer correction |  |  |  |  |
| --- | --- | --- | --- | --- |
|  | pValue | Observed Diff | Lower Limit | Upper Limit |
| HC – CE | 9.34e-5 | -3.30 | -4.68 | -1.93 |
| HC - BG | .001 | -2.47 | -3.84 | -1.09 |
| CE - BG | .28 | .84 | -.54 | 2.21 |

#### 3.1.2. Inter-Trial Phase coherence (ITPC)

While participants listened to isochronous auditory sequences presented at a *Sf* of 1.5Hz (Fig. 2A), their neural activity responded timely to tone onsets, as shown by the coherence peaks at 1.5Hz in the ITPC spectrum (Fig. 3A, left). A distinct peak at 1.5Hz was present in the ITPC spectrum for all groups. However, statistical analyses revealed significant group difference (Fig. 3A, right): HC (dark blue) showed stronger coherence at the *Sf* than both CE and BG groups (lighter shades of blue). The one-way ANOVA revealed a significant group effect (F(2,30) = 5.25, *p* = .01; Tab. 5), and post-hoc simple t-tests revealed higher coherence in HC than in both CE (*p* = .04) and BG (*p =* .02) groups, but no significant difference between CE and BG (Tab. 5; corrected for multiple comparisons via Tukey-Kramer). As the focus of this work is on delta-band neural dynamics centered at the stimulation frequency, we did not conduct any other statistical analysis on other frequency bands.

**Table 5.**
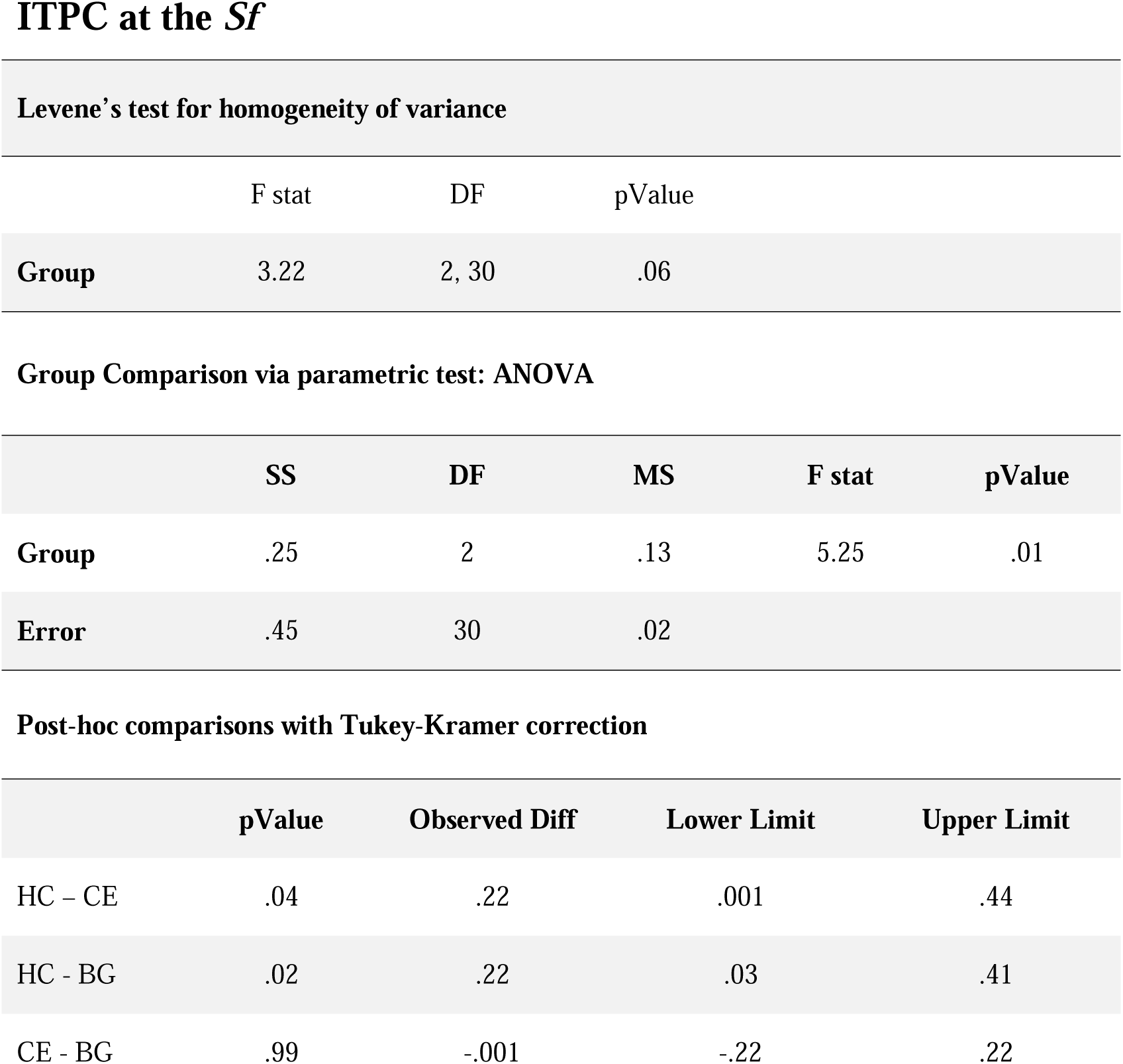
Group comparison of the ITPC at the stimulation frequency. In order, the table reports results from the Levene’s test for homogeneity of variance: F-statistics, degrees of freedom (DF) and p value. In the middle, the results from the ANOVA: sum of squares (SS), degrees of freedom (DF), mean squared error (MS), F-statistics and p-value. The error term reports the within-group variation. At the bottom, the results of the pairwise group comparison, with Tukey-Kramer correction. In order, the p-value, observed difference, lower and upper limit.

ITPC at the $S_f$
| Levene's test for homogeneity of variance |  |  |  |
| --- | --- | --- | --- |
|  | F stat | DF | pValue |
| **Group** | 3.22 | 2, 30 | .06 |

| Group Comparison via parametric test: ANOVA |  |  |  |  |  |
| --- | --- | --- | --- | --- | --- |
|  | SS | DF | MS | F stat | pValue |
| **Group** | .25 | 2 | .13 | 5.25 | .01 |
| **Error** | .45 | 30 | .02 |  |  |

Table 5 – Group comparison of the ITPC at the stimulation frequency.
| Post-hoc comparisons with Tukey-Kramer correction |  |  |  |  |
| --- | --- | --- | --- | --- |
|  | pValue | Observed Diff | Lower Limit | Upper Limit |
| HC – CE | .04 | .22 | .001 | .44 |
| HC - BG | .02 | .22 | .03 | .41 |
| CE - BG | .99 | -.001 | -.22 | .22 |

#### 3.1.3. Time-resolved ITPC

While ITPC and similar phase concentration measures are typically interpreted as a proxy of *entrainment*, it is generally hard to disentangle true entrainment from a sequela of evoked responses (Breaska & Deouell, 2017; Zoefel et al., 2018, Obleser et al., 2017; Haegens et al., 2017). In fact, simulations showed that evoked responses drive high phase coherence (Breska & Deouell, 2017; Obleser et al., 2017), ultimately weakening its functional link with temporal predictions. Thus, we estimated a time-resolved metric of ITPC (t-ITPC) to quantify the build-up of phase-coherence over the course of the auditory sequence. T-ITPC was obtained by quantifying phase coherence from the complex spectra of continuous wavelet transformed data in the delta frequency band. The group-level time course is displayed in Fig. 3B, color-coded per group. Next, we calculated, at the level of the single-participant, the slope of t-ITPC over a 5s interval starting from the third tone onset. Participant data were pooled into groups (Fig. 3B, right), and we later performed group comparisons. As the Levene’s test for homogeneity of variance was significant (F(2,30) = 3.34, *p* = .048; Tab 6), we performed non-parametric analysis of variance via a Kruskal-Wallis test. The Group effect was significant (Chi-sq(2,30) = 15.4, *p* < .001), and pairwise group comparisons performed via permutation testing (1000 permutations) revealed a significantly higher t-ITPC slope for HC than for both CE and BG patients (Tab. 6).

**Table 6.**
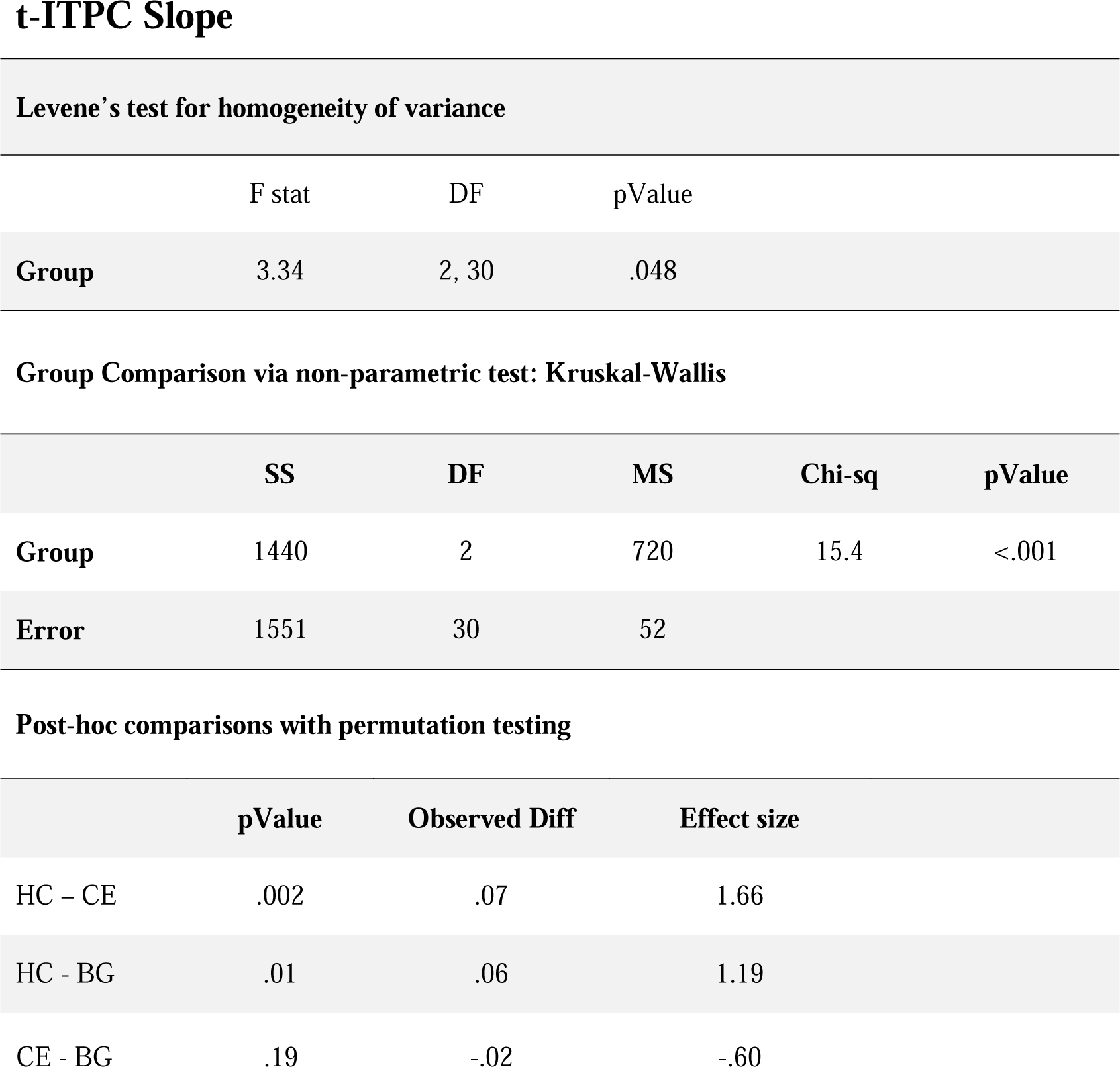
Group comparison of the t-ITPC Slope. In order, the table reports results from the Levene’s test for homogeneity of variance: F-statistics, degrees of freedom (DF) and p value. In the middle, the results from the Kruskal-Wallis test: sum of squares (SS), degrees of freedom (DF), mean squared error (MS), Chi-squared and p-value. The error term reports the within-group variation. At the bottom, the results of the pairwise group comparison performed via 1000 permutation tests. In order, the p-value, observed difference, effect size.

#### 3.1.4. Analyses on neural dynamics

##### Single-participant estimations

To assess the tuning of endogenous delta-band neural oscillatory activity toward the *Sf*, we quantified trial-level instantaneous frequency (IF), its dynamics of acceleration while tuning- and de-tuning to the *Sf*, its Stability (S), and Deviation (Dev) from the Sf. Each metric was estimated at the single-participant and –trial level as illustrated in Fig. 4A: on top, the time-course of delta-band filtered Hilbert-transformed data; in the middle, IF as the derivative of phase evolution over time, taking into account the sampling frequency; at the bottom, Acc was calculated as the derivative of IF. The dashed pink line on the IF plot indicates the *Sf*. On the right side, the distribution of IF and Acc in the ‘Pre-‘, ‘Post-’ and ‘Dur-‘(ing) the listening periods (color-coded in shades of blues), and pooling across the FC channel cluster. The time-series of IF and Acc were further used to estimate S, the stability of the neural signal. Finally, we obtained the Latency and Deviation (not visualized in the top panel): the first corresponds to the first crossing point of IF through the horizontal *Sf* line; the latter is the standard deviation from the *Sf*. Individual data were then pooled into 3 groups: HC, CE, and BG patients. The group-level data are provided in Fig. 4B and mirrors the layout of Fig. 4A, averaging across trials. On the right side, the violin plots show (in order) the IF and Acc, obtained by averaging data during the listening window (0-8s) and across channels and trials. Next to it, are the distributions of Dev, S and latency.

##### Group comparisons

We performed independent comparisons for the metrics mentioned above and focused on the listening period only (‘Dur’). The only group comparison showing a significant effect was for Stability: F(2,30) = 4.36, *p* = .02. Post-hoc simple tests showed that CE had lower Stability than HC (*p* = .016; Tab. 7), while no difference was found between HC-BG nor between CE-BG. The reduced Stability in CE patients reflects high variability in acceleration dynamics of delta-band activity, ultimately suggesting a poorer neural representation of temporal regularity displayed in the auditory sequences.

**Table 7.**
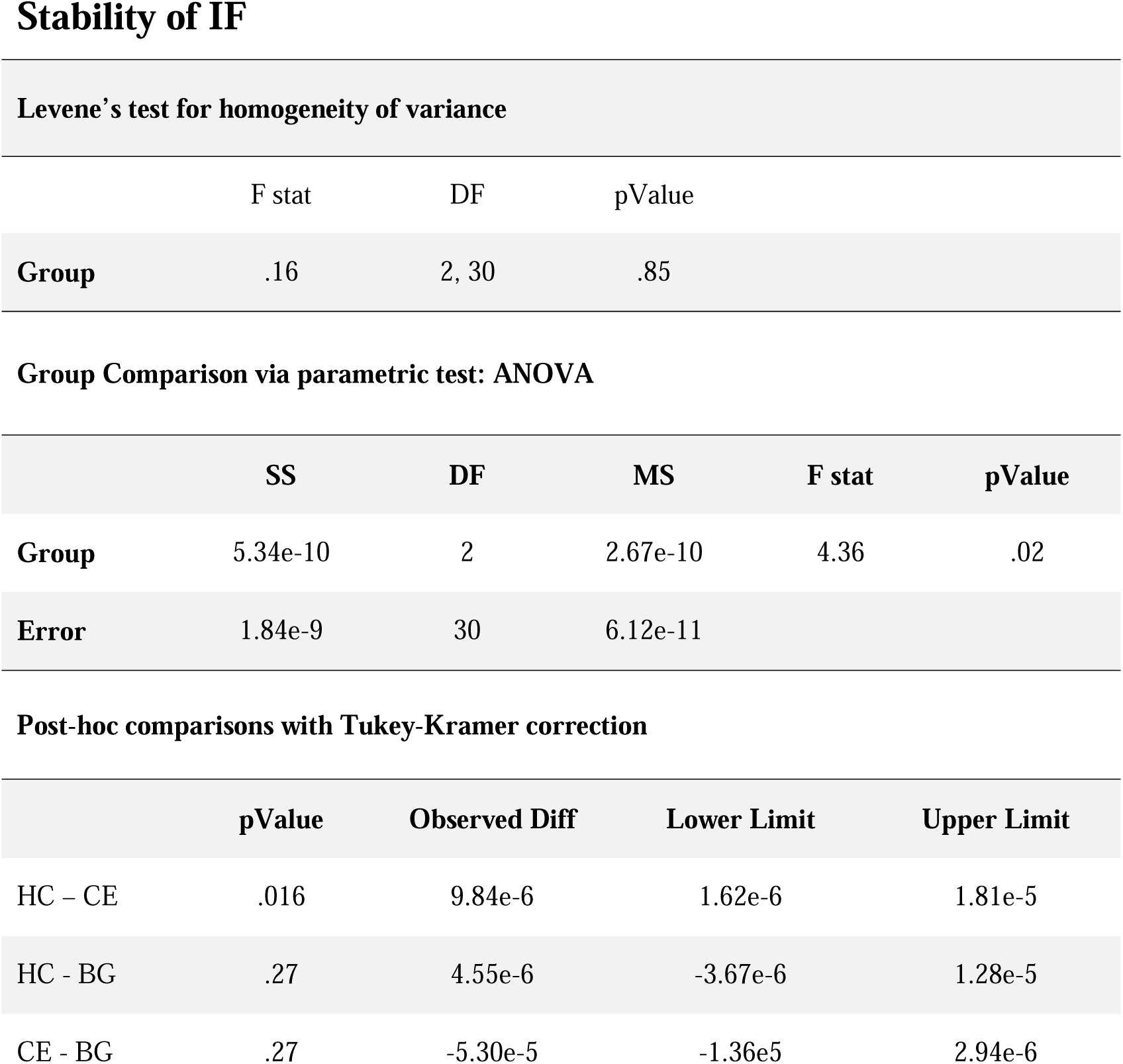
Group comparisons on the Stability of IF. In order, the table reports results from the Levene’s test for homogeneity of variance: F-statistics, degrees of freedom (DF) and p value. In the middle, the results from the ANOVA: sum of squares (SS), degrees of freedom (DF), mean squared error (MS), F-statistics and p-value. The error term reports the within-group variation. At the bottom, the results of the pairwise group comparison, with Tukey-Kramer correction. In order, the p-value, observed difference, lower and upper limit.

Levene's test for homogeneity of variance
|  | F stat | DF | pValue |
| --- | --- | --- | --- |
| **Group** | .16 | 2, 30 | .85 |

Group Comparison via parametric test: ANOVA
|  | SS | DF | MS | F stat | pValue |
| --- | --- | --- | --- | --- | --- |
| **Group** | 5.34e-10 | 2 | 2.67e-10 | 4.36 | .02 |
| **Error** | 1.84e-9 | 30 | 6.12e-11 |  |  |

Table 7 – Group comparisons on the Stability of IF.
|  | pValue | Observed Diff | Lower Limit | Upper Limit |
| --- | --- | --- | --- | --- |
| HC – CE | .016 | 9.84e-6 | 1.62e-6 | 1.81e-5 |
| HC - BG | .27 | 4.55e-6 | -3.67e-6 | 1.28e-5 |
| CE - BG | .27 | -5.30e-5 | -1.36e5 | 2.94e-6 |

### 3.2. Tapping experiment

Participants listened to auditory sequences at 3 rates (5, 10, 20Hz; Fig. 5A) and were asked to synchronize their tapping to the perceived beat and try to keep tapping at a comfortable rate. We assessed tapping behavior and variability over time by calculating inter-tap-intervals (ITI), its Shannon Entropy, converting it into frequency, and estimating Stability and Deviation from the 2.5Hz subharmonic. Group data are provided, in this order, in Fig. 5B. We excluded participants who did not complete the task and had less than 40 taps. As the resulting sample sizes did not allow any meaningful statistical comparisons, we will only derive qualitative observations in th discussion.

## 4. Discussion

We investigated the causal contributions of the cerebellum (CE) and the basal ganglia (BG) to sensory and sensorimotor processes underlying the detection, production and synchronization (‘DPS’) with basic auditory rhythms. Using EEG, we assessed the neural dynamics of temporal processing, unravelling group differences in *when, if,* and *how* healthy controls (HC), CE, and BG lesion patients process temporal regularities. Next, a behavioral experiment probed their capacity for DPS when tapping to temporally regular acoustic sequences at different rates.

First, we assessed the encoding of sound onset timing via event-related (ERP) and inter-trial phase coherence (ITPC) analyses. We found that CE patients showed lower N100 amplitudes, and increased N100 latency variability than HC. Along with a reduction of the ITPC at the stimulation frequency (*Sf*), these results suggest that CE lesions impacted the ability to process the precise timing of event onsets. In doing so, we support previous evidence documenting altered timing in CE patients (Grube, Cooper, et al., 2010; Nozaradan et al., 2017), typically leading to difficulties in producing stable rhythms, and synchronizing with external rhythms (Ivry & Keele, 1989; Ivry, Keele & Diener, 1988; Schwartze et al., 2016). However, the current results partially contradict other lines of evidence, showing intact sensitivity to the temporal structure of auditory sequences (Schwartze & Kotz, 2021; Breska & Ivry, 2018, 2020) in CE patients. In particular, CE lesions were found to causally impact the ability to estimate and process single intervals. However, the same patients were capable of estimating time intervals when embedded in a temporal structure (i.e., when the same interval is repeated a few times, leading to a rhythmic stream; Breska & Ivry, 2018, 2020, 2021). Consequently, CE patients should have benefited from the isochronous stimulation implemented here and should not have had difficulties in aligning their neural activity to tone onsets. Thus, were they not capable of generating and employing temporal predictions to guide their neural behavior? To assess this question, additional analyses investigated delta-band oscillations and their dynamics. We estimated a time-resolved metric of ITPC, aiming to quantify the build-up of phase-coherence over the course of the auditory sequence. Similar metrics were previously employed as an indicator of temporal predictions (Breska & Ivry, 2020) and auditory processing (Ten Oever et al., 2017). In fact, while ITPC peaks are typically driven by the phase rest of oscillations at tone onset (Breska & Deouell, 2017; Obleser et al., 2017), the t-ITPC slope could potentially reveal an underlying mechanism of increasing phase-alignment in anticipation of next sound onsets. CE patients’ t-ITPC slope was nearly flat, and significantly lower than HC, who showed an increasing slope over the course of the auditory sequence. This observation may indicate that CE patients were not capable of generating predictions about next tone onsets, as they were not coherently aligning delta-band oscillations over time. Were CE patients not capable of encoding a neural representation of temporal regularity? To address this question, we developed new metrics to quantify *tuning* dynamics of delta-band oscillations to the *Sf*. We calculated instantaneous frequency (IF), their dynamics of acceleration while encoding and tuning (to) (and de-tuning from) the *Sf*, and the stability of such neural representations. Results revealed a tendency toward an increased deviation from the *Sf*, and a significantly reduction in stability in CE compared to HC. The reduced stability might reflect high variability in acceleration dynamics in delta-band activity, ultimately suggesting altered neural encoding of the temporal regularity in isochronous sequences.

Altogether, increased variability in ERPs, reduced delta-band phase coherence, reduced slope of t-ITPC, and reduced stability of delta-band dynamics observed in CE patients suggest altered neural representation of temporal regularity in auditory sequences. CE lesions causally impacted the capacities to encode the precise timing of sound onsets, and further potentially impaired the ability to generate and employ temporal predictions to estimate next tone onsets. Finally, the data suggest that, differently from previous observations, CE patients did not benefit from temporal regularity in auditory sequences.

Moving to the BG lesion data, we found their ERPs’ amplitude and latency to be comparable to that of HC. However, similarly to CE patients, BG participants showed increased variability in the N100 peak amplitude latency. BG lesions were further associated with a significant reduction in ITPC peak at the *Sf*, and a reduced slope of the t-ITPC. These observations suggest altered encoding of temporal regularity in auditory sequences, supporting prior results (Nozaradan et al., 2017; Schwartze et al., 2015; Breska & Ivry, 2020). Furthermore, the reduction in the slope of the t-ITPC might suggest that BG lesions causally impacted the capacity to generate expectations about next tone onsets, thus aligning with and supporting prior evidence on a link between BG and temporal prediction (Grahn, 2009; Grahn & Brett, 2009a; Schwartze et al., 2011a; Teki, Grube, Kumar, et al., 2011).

However, the current results indicate striking similarities between CE and BG lesion patients: they did not show any significant difference in the employed neural metrics of temporal processing. These observations are at odds with prior evidence discussing fine, yet fundamental dissociations between rhythm- and interval-based temporal predictions in BG and CE lesion patients (e.g., Breska & Ivry, 2018; Schwartze & Kotz, 2013). Were their neural representations of the temporal regularities similar? Differently from CE, BG patients’ stability metric was comparable to HC, potentially indicating a more stable neural encoding of the *Sf*. Were these patients capable of employing compensatory mechanisms to overcome their difficulties in processing a basic rhythm? For instance, by estimating repeated temporal intervals? Previous literature showed that interval discrimination was deteriorated in rhythmic contexts for BG patients (Breska & Ivry, 2018), and large variability was observed in interval timing tasks at both the second and millisecond range (Jones et al., 2008; Merchant et al., 2008). Consequently, one may argue that BG patients should not be able to employ such a compensatory strategy. In fact, reduced ITPC, t-ITPC, and increased variability in the N100 latency all converge toward altered temporal processing. As such, the current results do now allow dissociating between a causal role of BG and CE. Why do the current results appear discrepant with the previous literature? At least two factors might have mediated these differences: the experimental paradigm and attention. In contrast to prior work, employing a few repeated temporal intervals and active task settings (e.g., Breska & Ivry, 2018), here we tested endogenous timing: participants were exposed for 20min to 8-to-11s long auditory sequences and their attention was diverted from the temporal aspects of the sequence as they were asked to count the N of deviant (amplitude attenuated) tones. Thus, the long exposure to an auditory isochronous stream and the absence of an explicit timing task allowed us to test temporal processing in absence of attention. Consequently, we might argue that, at least in passive listening, we cannot dissociate the roles of BG and CE. The use of isochronous auditory sequences, as opposed to complex auditory rhythms, might represent another limitation as these stimuli may have been suboptimal to dissociate CE-related sensory timing computations from BG-related temporal predictions. Finally, the small sample size might have not provided the adequate statistical power to discern such fine dissociations.

The tapping task might have helped in providing evidence for sensory and sensorimotor processes of temporal processing in these patients. However, most BG patients did not complete the task, and many participants across groups had a low number of taps. This observation suggests that the task was moderately complex, and particularly BG patients experienced difficulties in aligning their taps to the perceived temporal regularity. In fact, the task required participants to process the auditory sequence, extract the beat by detecting temporal regularity, and synchronize their motion to the external rhythm. In light of previous literature (e.g., Schwartze et al., 2015; Nozaradan et al., 2017), BG patients were expected to show altered tapping behaviors. The small sample size and the low statistical power, however, prevented us from interpreting these data meaningfully.

Overall, the newly employed methods allowed characterizing dynamics of delta-band oscillations in basic rhythm processing: we showed that delta oscillations fluctuate over time, *tuning in* and *out of* perceived acoustic regularity in auditory sequences by dynamically accelerating and slowing down from silence to listening periods, and further increasing phase coherence over time. These observations support the *dynamic attending* hypothesis (Jones & Boltz, 1989; Large & Jones, 1999), and confirm the role of low-frequency oscillations (Schroeder & Lakatos, 2009) in detecting, producing, and synchronizing (DPS) with temporal regularities in an auditory environment. On the other hand, the current approach also confirmed the fundamental role of the BG and CE in an extended cortico-subcortical-cortical network underlying rhythm and timing processing (Criscuolo, Pando-Naude, et al., 2022; Kasdan et al., 2022; Kotz et al., 2018; Schwartze & Kotz, 2013). In fact, we showed that while the healthy brain could flexibly and dynamically respond to and synchronize with sensory inputs, patients with lesions in the BG and CE did not. Patients showed a degree of heterogeneity and deteriorated capacity to synchronize ongoing neural activity to temporal regularities in the acoustic environment.

Differently from our hypotheses, however, our EEG data did not clearly dissociate CE from BG timing functions. We argued that our experimental setup and/or the adopted analytical approach may have played a role. Alternatively, it is possible that both patient populations employed compensatory mechanisms, thus masking out existing difference. In fact, we have previously discussed that BG and CE are part of a widespread cortico-subcortical network engaged in rhythm processing. Animal literature shows that there are neurons in the cortex acting as ‘neural chronometer’ encoding the passage of time and the precise timing of sensory events (Merchant et al., 2011; Merchant, Harrington, et al., 2013a; Merchant, Pérez, et al., 2013). These cortical neurons may take over timing computations as a functional reorganization mechanism after subcortical brain lesions, ultimately preventing to dissociate BG and CE’s roles in basic timing functions. Finally, another limitation lies in the difficulty to distinguish fine-grained timing processing in absence of spatio-temporally resolved data. In this perspective, this research field would benefit from a more accurate characterization of the time-course, strength and directionality of information flow during rhythm processing. Such approach would allow to monitor the interplay and the causal relationship between cortico-subcortical regions, as well as between the BG and CE. Furthermore, it would enable to characterize the frequency-specificity and directionality of influence of some timing computations. While we here focused on delta-band activity only, literature suggests beta-band activity to also be prominent in subcortical and cortical brain regions (Bartolo et al., 2014; Bartolo & Merchant, 2015; Keitel et al., 2017; Keitel & Gross, 2016) and to further be linked to predictive priming of sensory regions (Arnal, 2012; Engel & Fries, 2010). In other words, the anticipatory alignment of beta-band activity to the *when* of salient events (Fujioka et al., 2012, 2015) couples with bottom-up delta-band activity (Abbasi et al., 2018; Arnal et al., 2015; Merchant et al., 2015; Saleh et al., 2010) to instantiate motor-to-auditory predictions in support of adaptive behavior. Thus, our focus on scalp activity and on delta-band activity only may have prevented a full characterization of bottom-up and top-down mechanisms of temporal processing and prediction. Altogether, we encourage future studies to complement the current findings with investigations of beta-band activity, with a particular focus on event-related dynamics and beta-delta functional coupling. Such a complementary approach would deepen our understanding of the neurophysiological mechanisms underlying intra- and inter-individual variabilities in the capacities to detect, produce and synchronize with temporal regularities in the sensory environment, and to ultimately produce adaptive behavior.

## 5. Conclusions

The capacities to encode the precise timing of the sensory events around us, and to time our (re-)actions to changes in the environment are pivotal to act and adapt in a dynamically changing environment. In this study, we explored the rich and variegated landscape of neural oscillatory dynamics, and assessed *if, when* and *how* neural oscillations processed the temporal regularity in acoustic sequences. BG and CE lesions causally impacted the neurophysiological encoding of the rhythm, as demonstrated by reduced capacities to encode precise tone onsets and predict the timing next events.

## Author contributions

S.A.K., M.S., S.N., conceptualized both or one part of the study.

S.A.K., M.S., collected the data.

A.C. designed and performed data analyses.

A.C., S.A.K., M.S., S.N., interpreted the results and wrote the manuscript.

## Acknowledgments

The authors would like to thank Anne-Kathrin Franz, Christian Obermeier and Heike Boethel for support during data acquisition, Anika Stockert for support in the preparation of the clinical data and Julio Rodriguez-Larios for initial data discussion and analysis exploration. The authors would also like to thank the anonymous reviewers for their help in improving the overall quality of this article.

## Data and Code availability statement

The data and code in use here are available upon reasonable request to the corresponding author.

