## Supplementary Table for "Neural and behavioral dynamics of encoding, production and synchronization with external rhythms in subcortical lesion patients"

| Mixed effect model: ERP N100 Peak Amplitude Latency | | | | | | | | | | | | |
| --- | --- | --- | --- | --- | --- | --- | --- | --- | --- | --- | --- | --- |
| Model information: | | | | | | | | | | | | |
| Number of observations | | | 165 | | | | | | | | | |
| Fixed effects coefficients | | | 2 | | | | | | | | | |
| Random effects coefficients | | | 33 | | | | | | | | | |
| Covariance parameters | | | 2 | | | | | | | | | |
| Formula: | | | varOI ~ 1+ Group + (1 \| Participant) | | | | | | | | | |
| Model fit statistics | | | | | | | | | | | | |
| AIC | **BIC** | **Log Likelihood** | | | | | **Deviance** | |  | | | |
| -826 | -814 | 417.5 | | | | | -835 | |  | | | |
| Fixed effects coefficients (95% CI): | | | | | | | | | | | | |
|  | **Estimate** | **SE** | | **tStat** | | **DF** | | **pValue** | | **Lower** | | **Upper** |
| Intercept | .10 | .005 | | 19 | | 163 | | 3.67e-44 | | .09 | | .10 |
| Group | -.004 | .002 | | -.16 | | 163 | | .88 | | -.004 | | .004 |
| Random effects covariance parameters (95% CIs) | | | | | | | | | | | | |
|  | | **Type** | | | **Estimate** | | **Lower** | | **Upper** | |  | |
| Intercept \| Participant | | Std | | | .007 | | .004 | | .01 | |  | |
| Residual Std | | Std | | | .02 | | .016 | | .02 | |  | |

Suppl. Table 2 - Mixed effect model on ERP N100 Peak Amplitude Latency
The table reports model information: number of observations, fixed effect coefficients, random effect coefficients, covariance parameters. Then, the formula used to fit the model and model fit statistics: AIC, BIC values, Log Likelihood and Deviance. Further below, the fixed effect coefficients in a 95% confidence interval (CI): estimate, standard error, t-stat, degrees of freedom (DF), p value, lower and upper bound. Right below, random effects covariance parameters: estimate, lower and upper bound.
